# Retinal ganglion cell survival after severe optic nerve injury is modulated by crosstalk between JAK/STAT signaling and innate immune responses in the zebrafish retina

**DOI:** 10.1101/2021.04.08.439090

**Authors:** Si Chen, Kira L. Lathrop, Takaaki Kuwajima, Jeffrey M. Gross

## Abstract

Visual information is transmitted from the eye to the brain along the optic nerve, a structure composed of retinal ganglion cell (RGC) axons. The optic nerve is highly vulnerable to damage in neurodegenerative diseases like glaucoma and there are currently no FDA-approved drugs or therapies to protect RGCs from death. Zebrafish possess remarkable neuroprotective and regenerative abilities and here, utilizing an optic nerve transection (ONT) injury and an RNA-seq-based approach, we identify genes and pathways active in RGCs that may modulate their survival. Through pharmacological perturbation, we demonstrate that JAK/STAT pathway activity is required for RGC survival after ONT. Furthermore, we show that immune responses directly contribute to RGC death after ONT; macrophages/microglia are recruited to the retina and blocking neuroinflammation or depleting these cells after ONT rescues survival of RGCs. Taken together, our results support a model in which pro-survival signals in RGCs, mediated by JAK/STAT signaling, counteract the activity of innate immune responses to modulate RGC vulnerability and resilience in the zebrafish retina after severe optic nerve damage.

## INTRODUCTION

Visual information is transmitted from the eye to the brain along the optic nerve (ON), a structure composed of retinal ganglion cell (RGC) axons. The ON is highly vulnerable to damage and is compromised after acute injury and in neurodegenerative diseases such as glaucoma. In glaucoma, RGC axons are the initial site of injury; this causes the RGCs to die and ultimately results in irreversible loss of visual function. Neuroprotective strategies for glaucoma treatment seek to maintain the health of RGCs even after axons have been damaged, or to prevent initial damage to the RGC axon itself (Almasieh et al., 2012; Chang and Goldberg, 2012). There has been substantial progress in identifying the molecular and cellular events that lead to RGC death in the glaucomatous eye (Almasieh et al., 2012; Chang and Goldberg, 2012; Syc-Mazurek and Libby, 2019); however, no FDA-approved therapies currently exist to protect RGCs from death. This highlights the critical need for new neuroprotective strategies that preserve RGCs during glaucoma or after acute ocular trauma.

Most RGCs die in mammals suffering from glaucoma or acute ocular trauma. For example, in mouse, ∼65% of RGCs are lost within 7 days of optic nerve injury (ONI), and >90% by 28 days (Li et al., 2020). Mammals are also unable to regenerate RGCs after ONI, leading to irreparable vision loss. Unlike mammals, zebrafish possess remarkable neuroprotective and regenerative capacity in the central nervous system (Cigliola et al., 2020; Lahne et al., 2020). When the ON is damaged by crush or transection, zebrafish mount a robust regenerative response and regenerate RGC axons, restoring visual connections and function (Diekmann et al., 2015; Dhara et al., 2019). Moreover, it has been reported that ∼75% of zebrafish RGCs are protected from death after ONI, even to 7-weeks post-injury (Zou et al., 2013), but the mechanisms underlying neuroprotection are unknown. With an interest in developing novel strategies to preserve RGCs during glaucoma and other trauma, here, we identify potential neuroprotective factors/pathways in zebrafish that mediate RGC survival after ONI.

## MATERIALS AND METHODS

### Animals

Zebrafish (Danio rerio) in this study were 3-5 months old with an equal number of males and females used in all experiments. Transgenic lines used are *isl2b*:GFP (Pittman et al., 2008) and *mpeg1*:mCherry (Ellett et al., 2011); a gift from Dr. Neil Hukriede, University of Pittsburgh. Animals were maintained under standard conditions at 28.5 C on a 14h light/10 h dark cycle. There were no differences in outcomes based on gender of the fish and therefore all data were combined for analyses. All animals were treated in accordance with provisions established by the University of Pittsburgh School of Medicine Institutional Animal Care and Use Committee. Biological replicates (Ns) are provided in Figure legends for each experiment. At least three independent biological replicates were used per experiment.

### Optic nerve transection

Optic nerve transection (ONT) was performed as previously described (Elsaeidi et al., 2014; Zou et al., 2013). Zebrafish were anesthetized in 0.03% tricaine buffer (MS-222; Fisher Scientific) and placed on a moist tissue paper under a dissecting scope (Leica E65S). ONT surgery was performed on the left eye. After removal of the connective tissue, the eyeball was pulled out from the orbit gently using forceps. The ON and ophthalmic artery that runs along with the ON were exposed and the ON was then completely transected with another forcep, after which the eye was placed back in the orbit. Any animals where bleeding was observed were euthanized and not used for analysis. The right eye was subjected to a sham surgery as control: connective tissue was removed, the eye was pulled out from the orbit gently, and then placed back in the orbit. Fish were returned to system water in separate tanks to recover.

### RGC isolation and fluorescence-activated cell sorting (FACS)

Retinae were harvested from *isl2b*:GFP zebrafish at 12 and 24 hours post-injury (hpi) in biological triplicate. Four retinae were collected per sample. For retinal isolations and cell dissociation, animals were euthanized by tricaine overdose and transferred to PBS for enucleation. To achieve single cell suspension, the eyeball was rinsed in 1X PBS post-enucleation and incubated in StemPro™ Accutaset™ Cell Dissociation Reagent (Thermo Fisher, #A1110501) at 28.5 C for 40min in a water bath. The cell suspension was then passed through a 70μm cell strainer (Fisher Scientific) and gently pelleted by centrifugation at 4500rpm for 5min at 4C. After two washes in ice cold 1X PBS, cells were resuspended in ice cold 5% FBS in 1X PBS. GFP^+^ cells were sorted using a FACS Aria IIu cell sorter (BD Biosciences) at the Flow Cytometry Core at the University of Pittsburgh School of Medicine Department of Pediatrics. The gate for FACS was set by GFP intensity for both injured (ONT) and intact (control) *isl2b*:GFP samples. The same gating settings were used for both the ONT and control RGC samples and for all biological replicates.

### RNA-seq and bioinformatics analyses

Library preparation, quality control analysis, and next generation sequencing were performed by the Health Sciences Sequencing Core at Children’s Hospital of Pittsburgh as previously described (Leach et al., 2020). cDNA sequencing libraries were prepared using a SmartSeq HT kit (Takara Bio) and Illumina Nextera XT kit (Illumina Inc.). 2 ⨯ 75 paired-end, 150 cycle sequencing was performed on a NextSeq 500 system (Illumina Inc.), aiming for 40 million reads per sample. Raw read and processed data files are available in GEO: GSE171426. After sequencing, raw read data were imported to the CLC Genomics Workbench (Qiagen Digital Insights) licensed through the Molecular Biology Information Service of the Health Sciences Library System at the University of Pittsburgh. After mapping trimmed reads to the *Danio rerio* reference genome (assembly GRCz11), differentially expressed genes (DEG) from the 24hpi time point were identified using the following filter: the maximum of the average group RPKM value >1.5, absolute fold change >2, false discovery rate (FDR) p-value <0.05. Genes with TPM=0 in one or more replicates were excluded. This filtering strategy was also used for DEG at 12hpi and a second analysis was performed where the FDR p-value <0.05 was switched to a p-value <0.05. Pathway enrichment analyses were performed using DAVID Bioinformatics Resources 6.8 (https://david.ncifcrf.gov) and using the KEGG database.

### Pharmacological experiments

To assess toxicity and efficacy, the JAK inhibitor, Pyridone 6 (P6), or dexamethasone (both Sigma-Aldrich) was intravitreally (IV) injected into the intact and injured retina at several different concentrations, as previously described (Elsaeidi et al., 2014), and RGC survival was quantified. Based on (Elsaeidi et al. 2014; Bollaerts et al. 2019), 5uM P6 and 10uM dexamethasone were utilized. The first dose of each compound (2µL) was injected immediately after ONT (0dpi) and the second dose (2µL) was injected at 1dpi. 2μl of 0.05% DMSO was injected for all control doses. To deplete macrophages/microglia, fish were immersed in 500nM PLX3397 (Fisher Scientific) in system water, as previously described (Kanagaraj et al., 2020). PLX3397 exposure started 1 day before ONT and system water containing PLX3397 was replaced daily during the experiment.

### Immunohistochemistry

Immunofluorescence staining on retinal cryosections and flat-mounted retinae were performed as previously described (Uribe and Gross, 2007; Zou et al., 2013) with the addition of an antigen retrieval step consisting of 100% methanol incubation at −20 C for 30 minutes for staining pSTAT3 (MBL International Corporation, D128-3). For retinal flat-mounts, after euthanasia, fish were decapitated and heads were fixed in 4% paraformaldehyde (PFA) at 4C overnight. The retina was dissected in ice cold PBS, washed in 0.1% PBST (Triton X-100 in PBS), and then incubated in a 1.5 ml tube on a rotator overnight at 4C with 4C4 (1:200, a kind gift of Dr. Peter Hitchcock, University of Michigan School of Medicine; Craig et al., 2008), pSTAT3 (1:100, MBL), cleaved caspase 3 (1:200, Abcam, ab13847) and mCherry (1:200, Takara Bio USA Inc./Clontech Laboratories, 632543). Retinae were then washed in 0.1% PBST for 3 × 10 minutes at room temperature and incubated with goat-anti mouse Cy3 (1:250, Jackson ImmunoResearch Labs, 115-165-166) or goat-anti rabbit Alexa 647 (1:500, Cell Signaling Technology, 8940) secondary antibody for 3 hours at room temperature. Samples were then washed with 0.1% PBST for 3 × 10 minutes and carefully cut into 4 quadrants and mounted on slides with DAPI Vectashield (Vector Laboratories, H-1200). For cryosections, samples were prepared as previously described (Uribe and Gross, 2007); zn-8 (Zebrafish International Resource Center) was used at a 1:200 dilution and all other antibodies were used at the same concentrations as for the retinal flat-mount.

### BrdU Incorporation assays

Adult *isl2b*:GFP+ fish were immersed in 10mM BrdU (Sigma Aldrich) dissolved in system water for from 6dpi to 7dpi, and sacrificed at 7dpi. As a positive control for BrdU incorporation and immunohistochemistry, a needle poke injury was performed after (Fausett and Goldman, 2006) and fish were exposed to BrdU for 24 hours prior to being sacrificed. Immunohistochemistry for BrdU proceeded as described above for other antibodies, with the addition of a 10min incubation of 4N HCl at 37C to relax chromatin. anti-BrdU (Abcam, ab6326) was used at 1:200 dilution.

### Confocal microscopy, image processing and quantification

For *isl2b*:GFP imaging, retinal flat-mounts were prepared as above, with the head fixed in 4% PFA overnight at 4C and the retina dissected and mounted on the second day. Images were taken using Olympus Fluoview FV1200 laser scanning microscope (Olympus Corporation). Images were taken from each of the 4 quadrants (1 peripheral, 1 central per quadrant) at 40X magnification. Quantification of RGC numbers was performed using particle analysis in ImageJ after setting up a consistent threshold for all images. RGC survival was calculated as the ratio of *isl2b*:GFP^+^ RGCs in the left (ONT+) eye/ *isl2b*:GFP+ RGCs in the right (ONT- control) eye of the same fish.

Quantification of macrophages/microglia was performed using Imaris 9.6.0 (Bitplane). Confocal images were first converted into Imaris files, 3D rendered surfaces were then created for mCherry or 4C4 using the same algorithm (smoothing = 0.4μM, absolute intensity threshold = 1560, objects area > 50μM^2^) for each dataset. Quantification of total surface area and sphericity was performed using Imaris. Measurements were exported and statistically assessed using Prism 9.0 (Graphpad).

For GFP fluorescence intensity quantification, the GFP signal intensity of 30 random cells in the ONT+ and ONT- retina per fish (N=6) was collected using Fiji ImageJ and the corrected total cell fluorescence (CTCF) was obtained (after L. Hammond, 2014, *Measuring cell fluorescence using ImageJ*, The University of Queensland, Australia, https://theolb.readthedocs.io/en/latest/imaging/measuring-cell-fluorescence-using-imagej.html). Briefly, using the freehand tool in ImageJ, single RGCs were outlined as a region of interest (ROI) and GFP intensity was measured as was as the fluorescence of the background for every retina. CTCF was then calculated as follows: Integrated Density – (Area of selected cell X Mean fluorescence of background readings). Relative intensity was calculated as the ratio of the CTCF in ONT+ RGCs divided by that of ONT- RGCs.

Localization of pSTAT3 expression to RGCs was confirmed with surface creation in Imaris 9.6.0. Quantification of pSTAT3 levels was performed as described (Osborne et al. 2018) using FIJI. Briefly, *isl2b*:GFP labeling was used to identify the location and depth of the ganglion cell layer (GCL). RGC volumes were converted to Z-projections and background subtraction and speckle removal was performed on all images. Threshold levels for pSTAT3 were determined from ONT- samples and consistent thresholds were then applied to all images in ONT+ samples and integrated density was measured. The expression level of pSTAT3 was calculated as the ratio of the average integrated density of pSTAT3 in 1dpi, 7dpi, P6/1dpi, or P6/7dpi retinae to the average integrated density of the contralateral ONT- eye.

### Statistics

All statistical analyses were performed using Prism 9.0 (Graphpad). Data were presented as mean+/-SD, with the exception of cleaved caspase-3 images which show mean+/-SEM. For multiple comparisons, Kruskal-Wallis ANOVA followed by Dunn’s multiple comparisons tests between groups were performed. For comparisons between two groups, non-parametric Mann-Whitney tests were performed, with the exception of sphericity comparison in which an unpaired t-test with Welch’s correction was performed. P-values, sample sizes, and statistical analyses for each experiment are included in the figure legends.

## RESULTS AND DISCUSSION

### Zebrafish retain the majority of RGCs after ONT

To enable isolation of RGCs after injury, we first verified that GFP remained expressed in RGCs of adult *isl2b*:GFP fish (Pittman et al., 2008). In retinal cryosections, *isl2b*:GFP cells co-labeled with DAPI and the RGC-specific marker, zn-8 (Larison and Bremiller, 1990), and 64.86±8.44% of cells within the GCL were *isl2b*:GFP^+^ (Fig. 1A).

**Figure 1:**
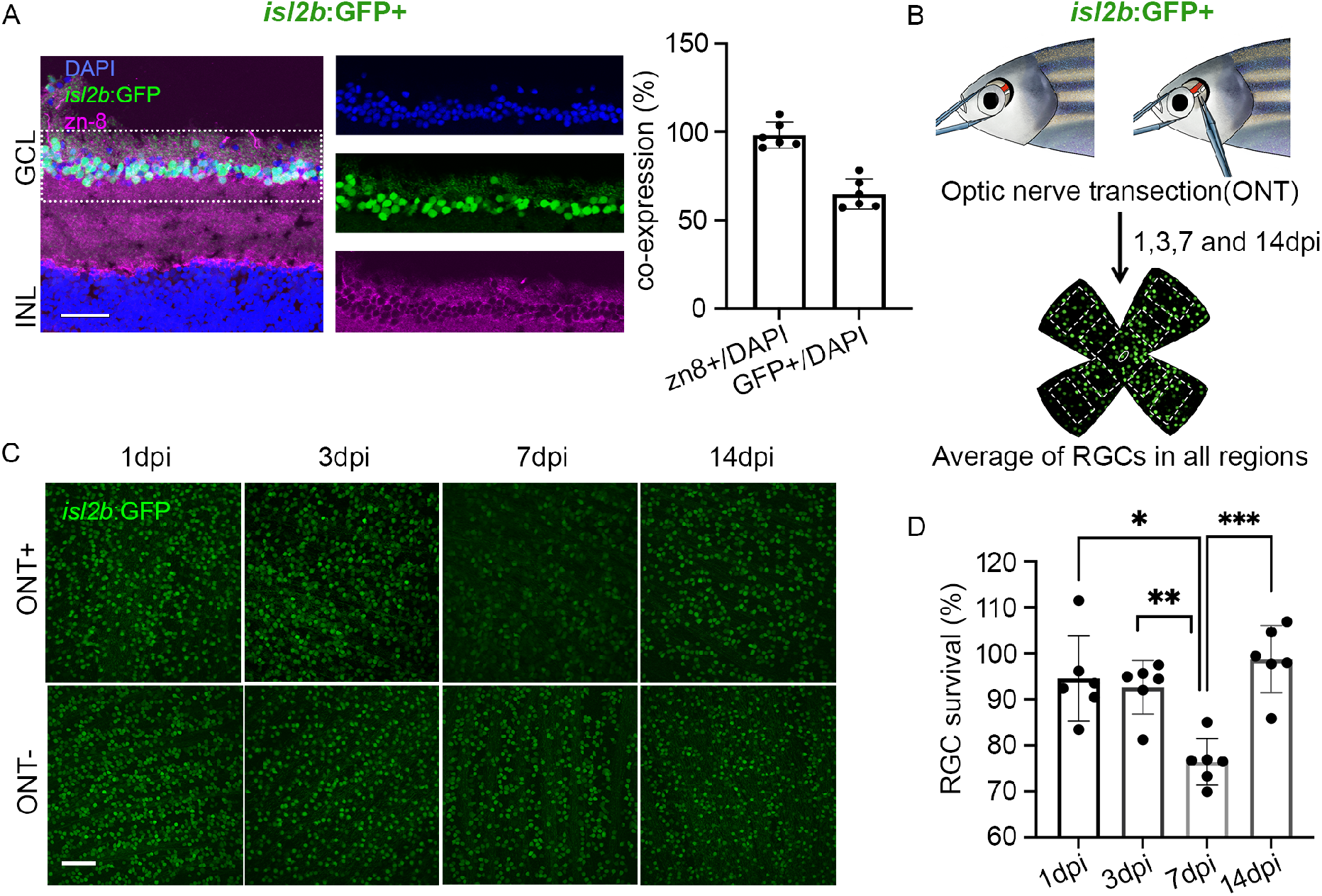
Zebrafish RGCs are preserved after ONT. **(A)** Immunolabeling of RGCs in the ganglion cell layer (GCL) with zn-8 (magenta) in the adult *isl2b*:GFP (green) retinae. ∼65% of DAPI (blue) stained RGCs were *isl2b*:GFP^+^. **(B)** Overview of optic nerve transection (ONT) and RGC survival analyses. **(C)** Images of 1, 3, 7, and 14dpi flat-mount retinae. **(D)** RGC survival percentages at 1, 3, 7, and 14dpi (n=6/day). Shown are mean±SD; *p<0.05; **p<0.01; ***p<0.001; Kruskal Wallis ANOVA w/ Dunn’s multiple comparisons. Scale bars = 50μm.

To create an ON injury, we performed optic nerve transection (ONT). Injured retinae from the left eye (ONT+) and sham surgery retinae from the right eye (ONT-) were collected 1, 3, 7, or 14 days post injury (dpi) (Fig. 1B). We confirmed that there were no differences in cell number between the uninjured left and right eye (S. Chen; *data not shown*). To quantify RGC survival after ONT, we counted *isl2b*:GFP^+^ RGCs from eight regions (four in the peripheral retina and four in the central retina (Fig. 1B)), then divided counts from the ONT+ retina by those from the ONT- retina of the same fish. RGC survival was 94.44±5.45% at 1dpi and 92.52±3.54% at 3dpi; however, survival decreased to 76.35±2.58% at 7dpi (p=0.0009). At 14dpi, RGC numbers recovered to 98.71±4.82% (Fig. 1C,D).

*isl2b*:GFP intensity in ONT+ RGCs declined by 40.43±9.99% at 7dpi relative to ONT- RGCs (p<0.0001), raising the possibility that GFP intensity was falling below the detection threshold after ONT due to compromised health and not death of *isl2b*:GFP^+^ RGCs. Cleaved caspase 3 immunostaining revealed that 12.44±1.79% of RGCs in the ONT+ retinae were caspase 3^+^ at 7dpi, compared to 0.06±0.01% of RGCs in the ONT- retina (Fig. S1). Moreover, no BrdU^+^ cells were detected in the GCL of ONT+ retinae at 7 dpi, indicating that RGCs had not yet regenerated (S. Chen, *data not shown)*. Taken together, these data indicate that, despite transient reduction in *isl2b*:GFP levels and limited death, most *isl2b*:GFP^+^ RGCs are indeed preserved in zebrafish after ONT.

### Identification of neuroprotective factors and pathways after ONT

To identify neuroprotective signals/pathways in RGCs, *isl2b*:GFP^+^ RGCs were FACS-isolated from ONT+ and ONT- retinae at 24hpi and utilized for RNA-seq (Fig 2A). We identified 308 differentially expressed genes (DEGs) (Fig. 2B), of which 56 were upregulated and 252 were downregulated (Tables S1,2). We reasoned that neuroprotective factors would be upregulated upon ONT, and the upregulated DEG group included *stat3, irf9, sox11b, lepr, and socs3b*, which encode components/regulators of the JAK/STAT signaling pathway (Fig. 2C; (Villarino et al., 2015)), in addition to other neuroprotective and pro-regenerative genes such as *gap43* (Chung et al., 2020), *atf3* (Kole et al., 2020), and *atoh7* (Brodie-Kommit et al., 2021). Moreover, the interferon regulatory factor genes, *irf9* and *irf1b* (Langevin et al., 2013), and the chemokine receptor, *cxcr4b* (García-Cuesta et al., 2019), were also upregulated, suggesting activation of innate immune responses in RGCs after ONT. Pathway enrichment analyses indicated that JAK/STAT signaling was the most highly enriched pathway in RGCs after ONT (Fig. 2D). Furthermore, DEGs associated with the adipocytokine signaling pathway, which regulates STAT3-mediated signals (Kadye et al., 2020), were also enriched in RGCs after ONT (Fig. 2D). Downregulated DEGs revealed that neuroactive ligand receptor interactions, MAPK, and calcium signaling pathways were all significantly changed after ONT (Fig. 2D).

**Figure 2:**
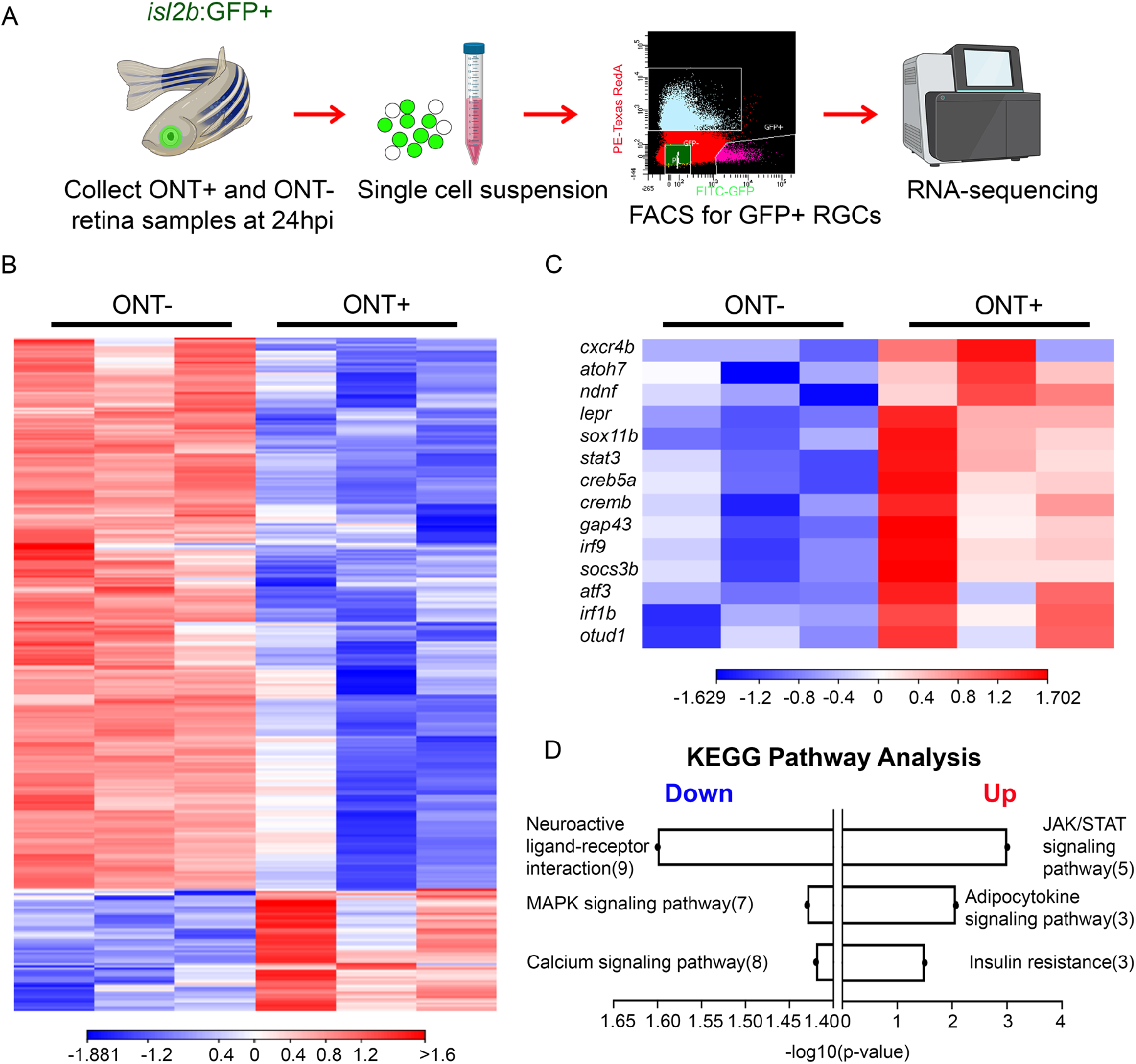
Identification of differentially expressed genes in *isl2b*:GFP^+^ RGCs after ONT. **(A)** Experimental workflow for FACS-isolation of *isl2b*:GFP^+^ RGCs. An example FACS plot showing a cell sorting gate is included. Icons were adapted from BioRender.com. **(B)** Heatmap showing hierarchical clustering of 308 DEGs at 24hpi from three biological replicates. **(C)** Heatmap highlighting DEGs of interest based on known neuroprotective and pro-regenerative functions. Heatmap legends show log_2_TPM. **(D)** Pathway enrichment analysis using the KEGG database showing top-3 down- and up-regulated pathways after ONT.

We also investigated gene expression changes at 12hpi. Using the same selection criteria as for 24hpi analyses yielded limited numbers of DEGs at 12hpi, with 5 upregulated and 3 downregulated (Table S3). Amongst these upregulated DEGs, however, was the leukocyte recruitment cytokine, *cxcl12b* (García-Cuesta et al., 2019), whose receptor, *cxcr4b*, was upregulated at 24hpi. Relaxing our selection criteria from an FDR p-value <0.05 to a p-value <0.05 revealed 49 upregulated and 31 downregulated DEGs (Tables S4,5). *stat3* and *socs3b* were amongst the upregulated group, as was the pro-inflammatory cytokine, *il1b* (Hasegawa et al., 2017), further suggesting activation of the JAK/STAT and innate immune pathways in RGCs after ONT.

### JAK/STAT pathway activity is required for RGC survival after ONT

JAK/STAT activity contributes to RGC survival in multiple injury contexts (Boyd et al., 2003; Huang et al., 2007; Luo et al., 2007) and facilitates ON and retinal regeneration (Elsaeidi et al., 2014; Kassen et al., 2009; Leibinger et al., 2013; Mehta et al., 2016; Park et al., 2004; Todd et al., 2016; Zhao et al., 2014). To confirm JAK/STAT pathway activation after ONT, we assessed phosphorylated-STAT3 (pSTAT3) levels in ONT+ and ONT- retinae (Fig. 3A; Movie S1). pSTAT3 levels were significantly increased in ONT+ RGCs (Fig. 3B; p<0.01), supporting the notion that Stat3 may be neuroprotective after ONT. To determine whether JAK/STAT pathway activity is required for RGC survival after ONT, we performed intravitreal (IV) injections of a pan-Jak inhibitor, Pyridone 6 (P6), or 0.05% DMSO at the time of ONT and again at 1dpi (Fig. 3C). P6-injection resulted in significant reductions to pSTAT3 levels in ONT+ RGCs at 1dpi and 7dpi, supporting the efficacy of P6 in blocking Jak activation in zebrafish (Fig. 3D,E). We next quantified *isl2b*:GFP^+^ RGC survival at 7dpi when Jak activity was impaired (Fig. 3F,G). DMSO had no effect on RGC survival, nor did P6 alone (p=0.4796; Fig. 3G). However, IV P6 injection in conjunction with ONT resulted in a significant reduction in RGC survival to 39.44±6.99% (p<0.0001). These data demonstrate a role for JAK/STAT pathway activity in protecting zebrafish RGCs after ONT.

**Figure 3:**
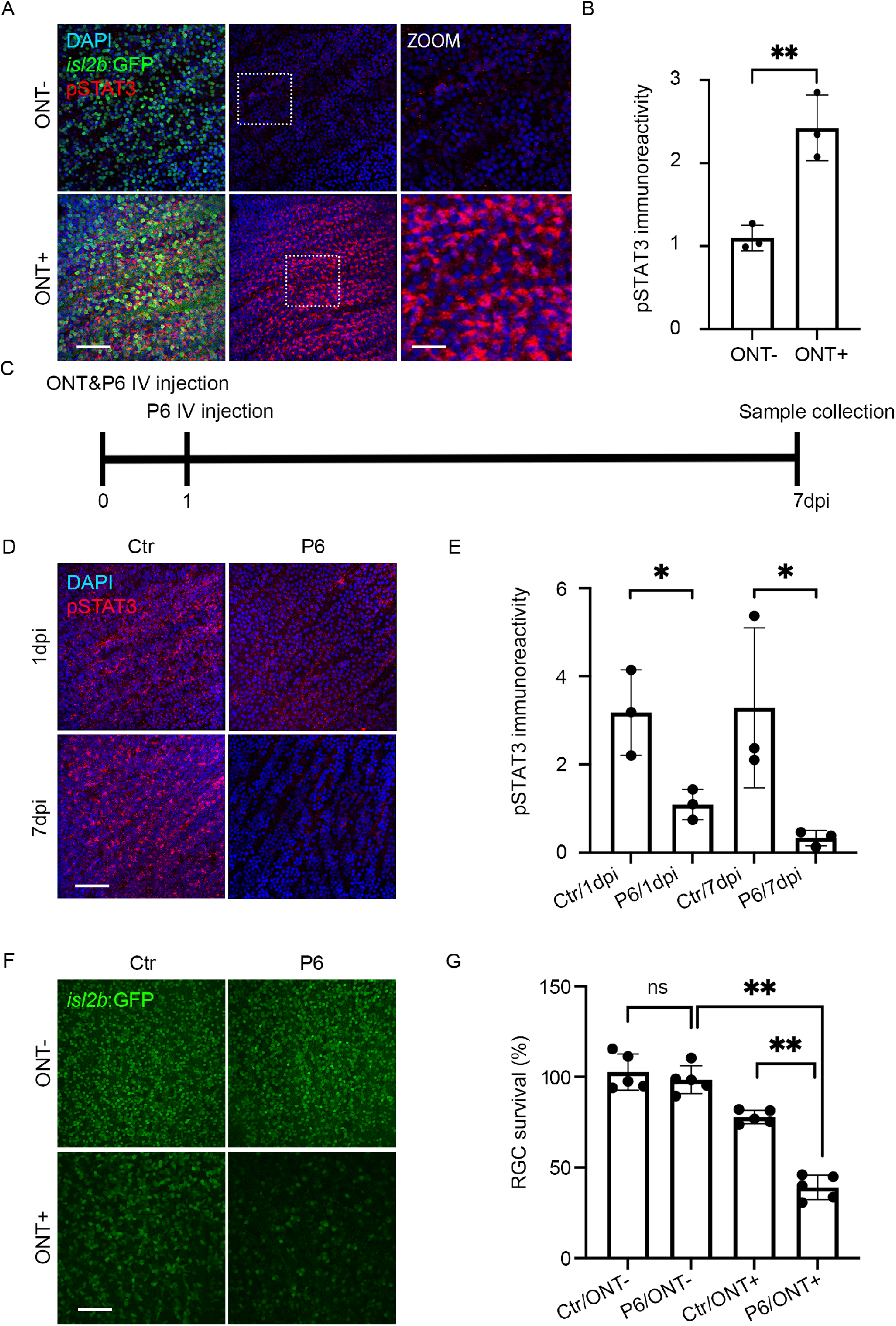
JAK/STAT pathway activity is required for RGC survival after ONT. **(A)** pSTAT3 expression (red) in flat-mount *isl2b*:GFP ONT- and ONT+ retinae at 1dpi. Nuclei stained with DAPI (blue). Boxed regions are 3x zooms of 40x images. **(B)** Quantification of pSTAT3 levels at 1dpi. pSTAT3 levels in ONT+ RGCs relative to levels normalized to those in ONT- RGCs. Shown are mean±SD of n=3 for each group; **p<0.01; Mann-Whitney test. **(C)** Experimental paradigm to assess Jak requirement during RGC survival after ONT. **(D)** pSTAT3 expression in ONT+ *isl2b*:GFP retinal flat mounts at 1 and 7dpi +/-intravitreal (IV) injection of the Jak inhibitor, P6. DMSO was used as control (Ctr). **(E)** Quantification of pSTAT3 expression after P6 application at 1 and 7dpi. N=3/condition. Shown are mean±SD; *p<0.05, Mann-Whitney test. **(F)** Images of 7dpi P6- or DMSO-injected flat-mount *isl2b*:GFP retinae. **(G)** Quantification of RGC survival after P6 injection (n=5/condition). Shown are mean±SD; **p<0.01; Kruskal Wallis ANOVA with Dunn’s multiple comparisons. Scale bars = 50μm and 150μm on zoomed images in A.

### Innate immune response involvement after ONT

After ONI, the effects of injury-activated immune responses are varied. In some contexts, immune responses stimulate the recruitment and activation of leukocytes that generate secondary signals to modulate RGC survival and death pathways (Bariş and Tezel, 2019; Mac Nair et al., 2016; Nadal-Nicolás et al., 2017; Williams et al., 2017), while in others, the stimulation of limited neuroinflammation induces pro-survival and regenerative responses (Kanagaraj et al., 2020; Todd et al., 2019). In zebrafish, leukocyte activity accelerates axonal regeneration after neuronal damage (Tsarouchas et al., 2018), including in RGCs after ON crush (Van Dyck et al., 2021). Immune response-related genes were upregulated in *isl2b*:GFP^+^ RGCs after ONT (Fig. 2) and therefore we investigated how components of the innate immune system respond to ONT in zebrafish and whether they contribute to RGC vulnerability. We focused on macrophages/microglia, leukocytes that accumulate in the zebrafish retina after a variety of injury types and facilitate repair and regeneration (Leach et al., 2020; White et al., 2017). In zebrafish, the 4C4 antibody recognizes an unidentified protein expressed by macrophages/microglia (Craig et al., 2008). Consistent with other reports (Mitchell et al., 2018), 4C4^+^ macrophages/microglia were located throughout the ONT- retina, including in the GCL (Fig. 4A,B; Movie S2). At 1dpi, the number of 4C4^+^ cells appeared to increase in the GCL (Fig. 4A,B). Similarly, utilizing *mpeg1*:mCherry transgenic fish (Ellett et al., 2011), we observed an apparent increase in mCherry^+^ macrophages/microglia in the ONT+ GCL at 1dpi (Fig. 4C). Due to morphology and close proximity in the ONT retinae, it was difficult to identify single macrophages/microglia for counting and therefore we quantified the percent area covered by mCherry^+^ cells within the GCL (Fig. 4D). When compared to ONT- retinae, mCherry^+^ cells covered nearly 6 times more GCL area after ONT (p=0.0006; Fig. 4D). Macrophage/microglia morphologies change upon activation, whereby quiescent cells with a ramified morphology take on a more spherical/amoeboid shape when activated (Karlstetter et al., 2015; Mitchell et al., 2018). Morphological differences were evident in both 4C4^+^ and mCherry^+^ cells within the GCL of ONT+ retinae (Fig. 4B,C), and quantification of mCherry^+^ cell sphericity revealed a significant increase at 1dpi (Fig. 4E; p<0.0001); collectively, showing that macrophages/microglia accumulate in the GCL and become activated after ONT.

**Figure 4:**
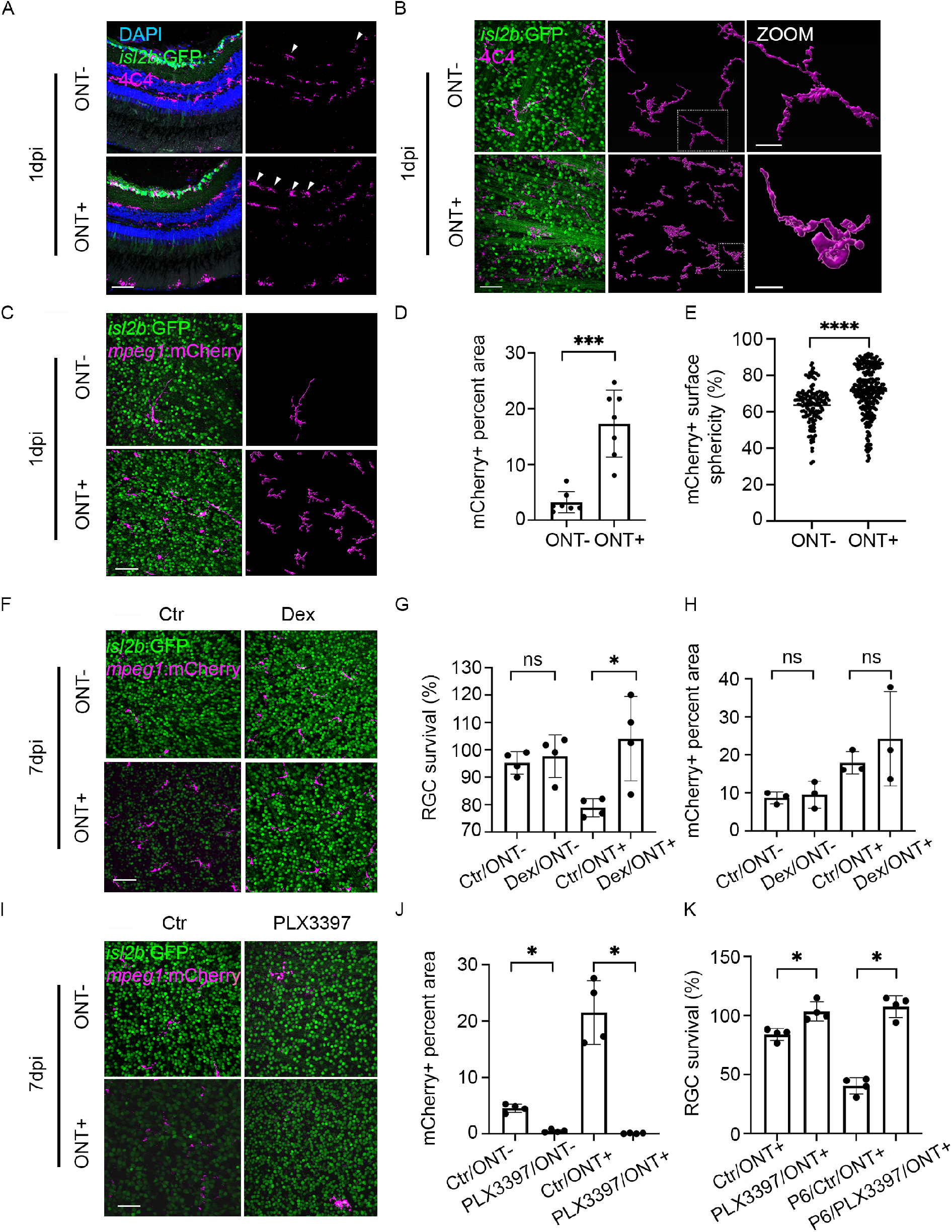
Macrophages/microglia are recruited to the GCL after ONT and mediate RGC death. Immunostaining of 4C4 (magenta) on *isl2b*:GFP retinal cryosections **(A)** and retinal flat mounts at 1dpi with Imaris surface renderings of 4C4^+^ macrophages/microglia **(B). (C)** Images of 1dpi retinal flat-mounts from *isl2b*:GFP;*mpeg1*:mCherry animals; macrophages/microglia (magenta). **(D)** Quantification of the GCL surface area occupied by mCherry^+^ macrophages/microglia at 1dpi (n=4/condition). Shown are mean±SD; ***p<0.001; Mann-Whitney test. **(E)** Violin plot showing a significant increase in sphericity of mCherry+ macrophages/microglia in ONT+ retinae compared to ONT- controls (n=140 in ONT- and n=272 in ONT+). ****p<0.0001; unpaired t-test with Welch’s correction. **(F)** Flat-mount images of *isl2b*:GFP;*mpeg1*:mCherry retinae after intravitreal injection of dexamethasone (Dex) or DMSO (Ctr) at 7dpi. **(G)** RGC survival in dexamethasone-treated retinae increased significantly at 7dpi when compared to control (n=4/condition). Shown are mean±SD; *p<0.05; Kruskal Wallis ANOVA test with Dunn’s multiple comparisons. **(H)** Quantification of mCherry^+^ macrophage/microglia coverage of the GCL after ONT and dexamethasone or DMSO injection (n=3/condition). Shown are mean±SD; Kruskal Wallis ANOVA test with Dunn’s multiple comparisons. No significant differences were detected. **(I)** Flat-mount images of *isl2b*:GFP;*mpeg1*:mCherry retinae after PLX3397 or control treatment (Ctr) at 7dpi. **(J)** Quantification of mCherry^+^ macrophage/microglia coverage of the GCL after ONT and PLX3397 treatment (n=4/condition). Shown are mean±SD; *p<0.05; Kruskal Wallis ANOVA test with Dunn’s multiple comparisons. **(K)** RGC survival in PLX3397-treated retinae increased significantly at 7dpi when compared to control. Similarly, RGC survival in PLX3397-treated retinae increased significantly after P6 addition over DMSO controls (n=4/condition). Shown are mean±SD; *P<0.05; Mann-Whitney test. Scale bars = 50μm.

### Blocking inflammation or depletion of macrophages/microglia protects RGCs after ONT

Pro-inflammatory cytokines, interferon response factors, and leukocyte recruitment factors were upregulated in *isl2b*:GFP^+^ RGCs post-ONT (Fig. 2C; Tables S1,S3,S4) and macrophages/microglia accumulate and become activated in the ONT+ GCL (Fig.4 A-D). These data suggest that macrophage/microglia-mediated inflammation might contribute to RGC death after ONT, in opposition to JAK/STAT-mediated neuroprotection. To test this hypothesis, we inhibited inflammation after ONT via dexamethasone treatment, a strategy shown to protect RGCs in other injury models (Bollaerts et al., 2019; Dutt et al., 2010; Gallina et al., 2015; Jovanovic et al., 2020), and quantified RGC survival. Experimentally, 2μL of 100μM dexamethasone or 0.05% DMSO (vehicle control) was IV injected into *isl2b*:GFP fish at the time of ONT and again at 1dpi and tissue was collected at 7dpi (after Fig. 3C). Dexamethasone alone had no effect on *isl2b*:GFP^+^ RGC survival in the ONT- retina (96.56±8.67%, p=0.5998), but significantly increased survival of ONT+ RGCs at 7dpi, compared to DMSO controls (104.1±13.35%, p<0.05; Fig. 4G). Dexamethasone-mediated inhibition of inflammation prevents the recruitment of macrophages/microglia in some contexts (Tsarouchas et al., 2018; White et al., 2017), but not others (Chatzopoulou et al., 2016; Warchol, 1999; Xie et al., 2019). Quantification of the percent of the GCL covered by *mpeg1*:mCherry^+^ macrophages/microglia in DMSO- and dexamethasone-injected retinae revealed no significant differences after ONT (Fig. 4H, p=0.0623). Taken together, these data demonstrate that impairment of inflammation after ONT rescued RGC survival but didn’t suppress the recruitment of macrophages/microglia to the GCL.

Previous reports have identified roles for microglia in contributing to RGC death after a variety of insults (Bosco et al., 2008; Jovanovic et al., 2020; Takeda et al., 2018). However, other studies have shown that microglia are dispensable for RGC survival after ON crush (Hilla et al., 2017), and may instead provide neuroprotective and/or pro-regenerative signals (Bell et al., 2018; Sappington et al., 2006). To directly test the requirement of macrophages/microglia in modulating RGC death in zebrafish after ONT, we depleted macrophages/microglia using PLX3397, a potent inhibitor of the colony-stimulating factor 1 receptor (CSF1R). CSF1R activity is required for macrophage/microglia differentiation (Lin et al., 2008; Sherr et al., 1985) and PLX3397 has been utilized effectively in zebrafish (e.g. Conedera et al., 2019; Leach et al., 2020; Van Dyck et al., 2021). Animals were immersed in 500nM PLX3397 one day prior to ONT and retinae were collected at 7dpi. To validate the efficiency of macrophage/microglia depletion by PLX3397, we quantified the percent area of the GCL occupied by *mpeg1*:mCherry^+^ macrophages/microglia. PLX3397 significantly reduced the coverage of *mpeg1*:mCherry^+^ cells in both the ONT- and ONT+ GCL (Fig. 4I,J). Similar to dexamethasone-mediated RGC protection (Fig. 4F,G), PLX3397-mediated depletion of macrophages/microglia also rescued RGC survival at 7dpi (103.49±12.01%, p<0.05; Fig. 4I,K). Finally, we determined whether the neuroprotective effects mediated by JAK/STAT activity in RGCs after ONT were required in the absence of macrophages/microglia. PLX3397-mediated depletion of macrophages/microglia also rescued RGC survival at 7dpi after JAK/STAT pathway inhibition using P6 (107.60±9.32%, p<0.05; Figs. 4K,S2), indicating that JAK/STAT activity is dispensable in the absence of macrophage/microglia recruitment.

Collectively, these data strongly support a model in which crosstalk between neurotoxic signals emanating from macrophages/microglia and JAK/STAT pathway activation in zebrafish RGCs regulates their survival after ONI. As noted above, Stat3 upregulation has been associated with RGC survival in some experimental contexts (Huang et al., 2007; Luo et al., 2007). Despite this, overall RGC survival is limited under physiological conditions, with over 90% of RGCs lost within 28 days after injury in mice (Li et al., 2020). This is not true for all mammals, however. Indeed, the naked mole-rat retains many RGCs after injury, to at least 28 days (Park et al., 2017). Interestingly, while pStat3 is nearly absent in mouse RGCs, even after injury, it increases significantly in mole-rat RGCs after ON crush, supporting a possible role for Stat3 activity in RGC neuroprotection.

Counter to our expectations, pSTAT3 localization was predominantly cytoplasmic in ONT+ RGCs, rather than nuclear (Fig. 3A, Movie S1). Interestingly, this observation is consistent with cytoplasmic pSTAT3 localization in the regenerating zebrafish retina (Elsaeidi et al., 2014) and in mouse motor neurons responding to cytokine stimulation (Selvaraj et al., 2012). Stat3 possesses transcription-independent functions such as cytoplasmically regulating autophagy (Shen et al., 2012). Stat3 also localizes to mitochondria after cytokine stimulation in mouse RGCs where it regulates metabolic functions and enhances axon regeneration after ONI (Luo et al., 2016). Thus, it is possible that the function of Stat3 in RGC neuroprotection could be transcription-independent.

Not all zebrafish RGCs survive ONT and caspase 3^+^ RGCs appeared to be a non-random pattern in the ONT+ retina (Fig. S1). This may indicate that there are RGC subtype(s) that are more susceptible to ONI, similar to what has been observed in mice (Tran et al., 2019). RGC subtypes in zebrafish have been recently characterized (Kölsch et al., 2021) and additional studies will be required to determine if specific subtypes are lost after ONT, and if so, whether these subtypes lack the ability to upregulate JAK/STAT activity after injury. Finally, it will be of interest to identify the signals emanating from macrophages/microglia that activate death and/or pro-survival pathways in zebrafish RGCs after injury, as these would also be promising targets around which neuroprotective therapies for glaucoma could be developed (García-Bermúdez et al., 2021; Rashid et al., 2019).

## ACKNOWLEDGEMENTS

The work described herein was supported by a grant from the BrightFocus Foundation National Glaucoma Research Program (G2020277) to JMG; an unrestricted grant from Xiangya Hospital of Central South University and China Scholar Council for studying in Pittsburgh to SC; NIH CORE Grant P30-EY08098 to the Department of Ophthalmology; the Eye & Ear Foundation of Pittsburgh, and an unrestricted grant from Research to Prevent Blindness, New York, NY. We’re grateful to Dick Barrett and Ben Carr for technical assistance and to Dr. Hugh Hammer for expert zebrafish husbandry.

## LITERATURE CITED

Almasieh, M., Wilson, A. M., Morquette, B., Cueva Vargas, J. L. and Di Polo, A. (2012). The molecular basis of retinal ganglion cell death in glaucoma. Prog. Retin. Eye Res. 31, 152–181.

Bariş, M. and Tezel, G. (2019). Immunomodulation as a Neuroprotective Strategy for Glaucoma Treatment. Curr Ophthalmol Rep 7, 160–169.

Bell, K., Und Hohenstein-Blaul, N. von T., Teister, J. and Grus, F. (2018). Modulation of the Immune System for the Treatment of Glaucoma. Curr. Neuropharmacol. 16, 942–958.

Bollaerts, I., Van Houcke, J., Beckers, A., Lemmens, K., Vanhunsel, S., De Groef, L. and Moons, L. (2019). Prior Exposure to Immunosuppressors Sensitizes Retinal Microglia and Accelerates Optic Nerve Regeneration in Zebrafish. Mediators Inflamm. 2019 6135795. https://doi.org/10.1155/2019/6135795

Bosco, A., Inman, D. M., Steele, M. R., Wu, G., Soto, I., Marsh-Armstrong, N., Hubbard, W. C., Calkins, D. J., Horner, P. J. and Vetter, M. L. (2008). Reduced retina microglial activation and improved optic nerve integrity with minocycline treatment in the DBA/2J mouse model of glaucoma. Invest. Ophthalmol. Vis. Sci. 49, 1437–1446.

Boyd, Z. S., Kriatchko, A., Yang, J., Agarwal, N., Wax, M. B. and Patil, R. V. (2003). Interleukin-10 Receptor Signaling through STAT-3 Regulates the Apoptosis of Retinal Ganglion Cells in Response to Stress. Investigative Ophthalmology & Visual Science 44, 5206–5211.

Brodie-Kommit, J., Clark, B. S., Shi, Q., Shiau, F., Kim, D. W., Langel, J., Sheely, C., Ruzycki, P. A., Fries, M., Javed, A., et al. (2021). Atoh7-independent specification of retinal ganglion cell identity. Sci Adv 7, 11 eabe4983.

Chang, E. E. and Goldberg, J. L. (2012). Glaucoma 2.0: neuroprotection, neuroregeneration, neuroenhancement. Ophthalmology 119, 979–986.

Chatzopoulou, A., Heijmans, J. P. M., Burgerhout, E., Oskam, N., Spaink, H. P., Meijer, A. H. and Schaaf, M. J. M. (2016). Glucocorticoid-Induced Attenuation of the Inflammatory Response in Zebrafish. Endocrinology 157, 2772–2784.

Chung, D., Shum, A. and Caraveo, G. (2020). GAP-43 and BASP1 in Axon Regeneration: Implications for the Treatment of Neurodegenerative Diseases. Front Cell Dev Biol 8, doi: 10.3389/fcell.2020.567537.

Cigliola, V., Becker, C. J. and Poss, K. D. (2020). Building bridges, not walls: spinal cord regeneration in zebrafish. Dis. Model. Mech. 13, doi: 10.1242/dmm.044131

Conedera, F. M., Pousa, A. M. Q., Mercader, N., Tschopp, M. and Enzmann, V. (2019). Retinal microglia signaling affects Müller cell behavior in the zebrafish following laser injury induction. Glia 67, 1150–1166.

Craig, S. E. L., Calinescu, A.-A. and Hitchcock, P. F. (2008). Identification of the molecular signatures integral to regenerating photoreceptors in the retina of the zebrafish. J. Ocul. Biol. Dis. Infor. 1, 73–84.

Dhara, S. P., Rau, A., Flister, M. J., Recka, N. M., Laiosa, M. D., Auer, P. L. and Udvadia, A. J. (2019). Cellular reprogramming for successful CNS axon regeneration is driven by a temporally changing cast of transcription factors. Sci. Rep. 9, 14198. doi: 10.1038/s41598-019-50485-6.

Diekmann, H., Kalbhen, P. and Fischer, D. (2015). Characterization of optic nerve regeneration using transgenic zebrafish. Front. Cell. Neurosci. 9, 118. doi: 10.3389/fncel.2015.00118.

Dutt, M., Tabuena, P., Ventura, E., Rostami, A. and Shindler, K. S. (2010). Timing of corticosteroid therapy is critical to prevent retinal ganglion cell loss in experimental optic neuritis. Investigative Ophthalmology & Visual Science 51, 1439–1445.

Ellett, F., Pase, L., Hayman, J. W., Andrianopoulos, A. and Lieschke, G. J. (2011). mpeg1 promoter transgenes direct macrophage-lineage expression in zebrafish. Blood 117, e49–56.

Elsaeidi, F., Bemben, M. A., Zhao, X.-F. and Goldman, D. (2014). Jak/Stat signaling stimulates zebrafish optic nerve regeneration and overcomes the inhibitory actions of Socs3 and Sfpq. J. Neurosci. 34, 2632–2644.

Fausett, B. V. and Goldman, D. (2006). A Role for α1 Tubulin-Expressing Müller Glia in Regeneration of the Injured Zebrafish Retina. J. Neurosci. 26, 6303–6313.

Gallina, D., Zelinka, C. P., Cebulla, C. M. and Fischer, A. J. (2015). Activation of glucocorticoid receptors in Müller glia is protective to retinal neurons and suppresses microglial reactivity. Exp. Neurol. 273, 114–125.

García-Bermúdez, M. Y., Freude, K. K., Mouhammad, Z. A., van Wijngaarden, P., Martin, K. K. and Kolko, M. (2021). Glial Cells in Glaucoma: Friends, Foes, and Potential Therapeutic Targets. Front. Neurol. 12, doi: 10.3389/fneur.2021.624983.

García-Cuesta, E. M., Santiago, C. A., Vallejo-Díaz, J., Juarranz, Y., Rodríguez-Frade, J. M. and Mellado, M. (2019). The Role of the CXCL12/CXCR4/ACKR3 Axis in Autoimmune Diseases. Front. Endocrinol. 10, doi: 10.3389/fendo.2019.00585.

Hasegawa, T., Hall, C. J., Crosier, P. S., Abe, G., Kawakami, K., Kudo, A. and Kawakami, A. (2017). Transient inflammatory response mediated by interleukin-1β is required for proper regeneration in zebrafish fin fold. Elife 6, https://doi.org/10.7554/eLife.22716.

Hilla, A. M., Diekmann, H. and Fischer, D. (2017). Microglia Are Irrelevant for Neuronal Degeneration and Axon Regeneration after Acute Injury. J. Neurosci. 37, 6113–6124.

Huang, Y., Cen, L.-P., Choy, K. W., van Rooijen, N., Wang, N., Pang, C. P. and Cui, Q. (2007). JAK/STAT pathway mediates retinal ganglion cell survival after acute ocular hypertension but not under normal conditions. Exp. Eye Res. 85, 684–695.

Huang, T., Li, H., Zhang, S., Liu, F., Wang, D. and Xu, J. (2021). Nrn1 Overexpression Attenuates Retinal Ganglion Cell Apoptosis, Promotes Axonal Regeneration, and Improves Visual Function Following Optic Nerve Crush in Rats. Journal of Molecular Neuroscience 71, 66–79.

Jovanovic, J., Liu, X., Kokona, D., Zinkernagel, M. S. and Ebneter, A. (2020). Inhibition of inflammatory cells delays retinal degeneration in experimental retinal vein occlusion in mice. Glia 68, 574–588.

Kadye, R., Stoffels, M., Fanucci, S., Mbanxa, S. and Prinsloo, E. (2020). A STAT3 of Addiction: Adipose Tissue, Adipocytokine Signalling and STAT3 as Mediators of Metabolic Remodelling in the Tumour Microenvironment. Cells 9, 1043. doi:10.3390/cells9041043.

Kanagaraj, P., Chen, J. Y., Skaggs, K., Qadeer, Y., Conner, M., Cutler, N., Richmond, J., Kommidi, V., Poles, A., Affrunti, D., et al. (2020). Microglia Stimulate Zebrafish Brain Repair Via a Specific Inflammatory Cascade. Cold Spring Harbor Laboratory 2020.10.08.330662.

Karlstetter, M., Scholz, R., Rutar, M., Wong, W. T., Provis, J. M. and Langmann, T. (2015). Retinal microglia: just bystander or target for therapy? Prog. Retin. Eye Res. 45, 30–57.

Kassen, S. C., Thummel, R., Campochiaro, L. A., Harding, M. J., Bennett, N. A. and Hyde, D. R. (2009). CNTF induces photoreceptor neuroprotection and Müller glial cell proliferation through two different signaling pathways in the adult zebrafish retina. Exp. Eye Res. 88, 1051–1064.

Kole, C., Brommer, B., Nakaya, N., Sengupta, M., Bonet-Ponce, L., Zhao, T., Wang, C., Li, W., He, Z. and Tomarev, S. (2020). Activating Transcription Factor 3 (ATF3) Protects Retinal Ganglion Cells and Promotes Functional Preservation After Optic Nerve Crush. Invest. Ophthalmol. Vis. Sci. 61, 31. doi:10.1167/iovs.61.2.31

Kölsch, Y., Hahn, J., Sappington, A., Stemmer, M., Fernandes, A. M., Helmbrecht, T. O., Lele, S., Butrus, S., Laurell, E., Arnold-Ammer, I., et al. (2021). Molecular classification of zebrafish retinal ganglion cells links genes to cell types to behavior. Neuron 109, 645–662.e9.

Lahne, M., Nagashima, M., Hyde, D. R. and Hitchcock, P. F. (2020). Reprogramming Müller Glia to Regenerate Retinal Neurons. Annu Rev Vis Sci 6, 171–193.

Langevin, C., Aleksejeva, E., Passoni, G., Palha, N., Levraud, J.-P. and Boudinot, P. (2013). The antiviral innate immune response in fish: evolution and conservation of the IFN system. J. Mol. Biol. 425, 4904–4920.

Larison, K. D. and Bremiller, R. (1990). Early onset of phenotype and cell patterning in the embryonic zebrafish retina. Development 109, 567–576.

Leach, L. L., Hanovice, N. J., George, S. M., Gabriel, A. E. and Gross, J. M. (2020). The immune response is a critical regulator of zebrafish retinal pigment epithelium regeneration. biorxiv 2020.08.14.250043.

Leibinger, M., Andreadaki, A., Diekmann, H. and Fischer, D. (2013). Neuronal STAT3 activation is essential for CNTF-and inflammatory stimulation-induced CNS axon regeneration. Cell Death & Disease 4, e805–e805.

Li, L., Huang, H., Fang, F., Liu, L., Sun, Y. and Hu, Y. (2020). Longitudinal Morphological and Functional Assessment of RGC Neurodegeneration After Optic Nerve Crush in Mouse. Front. Cell. Neurosci. 14, 109. doi:10.3389/fncel.2020.00109

Lin, H., Lee, E., Hestir, K., Leo, C., Huang, M., Bosch, E., Halenbeck, R., Wu, G., Zhou, A., Behrens, D., et al. (2008). Discovery of a cytokine and its receptor by functional screening of the extracellular proteome. Science 320, 807–811.

Luo, J.-M., Cen, L.-P., Zhang, X.-M., Chiang, S. W.-Y., Huang, Y., Lin, D., Fan, Y.-M., Van Rooijen, N., Lam, D. S. C., Pang, C. P., et al. (2007). PI3K/akt, JAK/STAT and MEK/ERK pathway inhibition protects retinal ganglion cells via different mechanisms after optic nerve injury. European Journal of Neuroscience 26, 828–842.

Luo, X., Ribeiro, M., Bray, E. R., Lee, D.-H., Yungher, B. J., Mehta, S. T., Thakor, K. A., Diaz, F., Lee, J. K., Moraes, C. T., et al. (2016). Enhanced Transcriptional Activity and Mitochondrial Localization of STAT3 Co-induce Axon Regrowth in the Adult Central Nervous System. Cell Rep. 15, 398–410.

Mac Nair, C. E., Schlamp, C. L., Montgomery, A. D., Shestopalov, V. I. and Nickells, R. W. (2016). Retinal glial responses to optic nerve crush are attenuated in Bax-deficient mice and modulated by purinergic signaling pathways. J. Neuroinflammation 13, 93. https://doi.org/10.1186/s12974-016-0558-y

Mehta, S. T., Luo, X., Park, K. K., Bixby, J. L. and Lemmon, V. P. (2016). Hyperactivated Stat3 boosts axon regeneration in the CNS. Exp. Neurol. 280, 115–120.

Mitchell, D. M., Lovel, A. G. and Stenkamp, D. L. (2018). Dynamic changes in microglial and macrophage characteristics during degeneration and regeneration of the zebrafish retina. J. Neuroinflammation 15, 163. https://doi.org/10.1186/s12974-018-1185-6

Nadal-Nicolás, F. M., Jiménez-López, M., Salinas-Navarro, M., Sobrado-Calvo, P., Vidal-Sanz, M. and Agudo-Barriuso, M. (2017). Microglial dynamics after axotomy-induced retinal ganglion cell death. J. Neuroinflammation 14, 218. https://doi.org/10.1186/s12974-017-0982-7

Osborne, A., Khatib, T. Z., Songra, L., Barber, A. C., Hall, K., Kong, G. Y. X., Widdowson, P. S. and Martin, K. R. (2018). Neuroprotection of retinal ganglion cells by a novel gene therapy construct that achieves sustained enhancement of brain-derived neurotrophic factor/tropomyosin-related kinase receptor-B signaling. Cell Death Dis. 9, 1007. https://doi.org/10.1038/s41419-018-1041-8

Park, K. K., Luo, X., Mooney, S. J., Yungher, B. J., Belin, S., Wang, C., Holmes, M. M. and He, Z. (2017). Retinal ganglion cell survival and axon regeneration after optic nerve injury in naked mole-rats. J. Comp. Neurol. 525, 380–388.

Park, K., Luo, J.-M., Hisheh, S., Harvey, A. R. and Cui, Q. (2004). Cellular mechanisms associated with spontaneous and ciliary neurotrophic factor-cAMP-induced survival and axonal regeneration of adult retinal ganglion cells. J. Neurosci. 24, 10806–10815.

Pereiro, X., Ruzafa, N., Acera, A., Fonollosa, A., Rodriguez, F. D. and Vecino, E. (2018). Dexamethasone protects retinal ganglion cells but not Müller glia against hyperglycemia in vitro. PLoS One 13, e0207913.

Pittman, A. J., Law, M.-Y. and Chien, C.-B. (2008). Pathfinding in a large vertebrate axon tract: isotypic interactions guide retinotectal axons at multiple choice points. Development 135, 2865–2871.

Rashid, K., Akhtar-Schaefer, I. and Langmann, T. (2019). Microglia in Retinal Degeneration. Front. Immunol. 10, 1975. doi: 10.3389/fimmu.2019.01975.

Sappington, R. M., Chan, M. and Calkins, D. J. (2006). Interleukin-6 protects retinal ganglion cells from pressure-induced death. Invest. Ophthalmol. Vis. Sci. 47, 2932–2942.

Selvaraj, B. T., Frank, N., Bender, F. L. P., Asan, E. and Sendtner, M. (2012). Local axonal function of STAT3 rescues axon degeneration in the pmn model of motoneuron disease. J. Cell Biol. 199, 437–451.

Shen, S., Niso-Santano, M., Adjemian, S., Takehara, T., Malik, S. A., Minoux, H., Souquere, S., Mariño, G., Lachkar, S., Senovilla, L., et al. (2012). Cytoplasmic STAT3 represses autophagy by inhibiting PKR activity. Mol. Cell 48, 667–680.

Sherr, C. J., Rettenmier, C. W., Sacca, R., Roussel, M. F., Look, A. T. and Stanley, E. R. (1985). The c-fms proto-oncogene product is related to the receptor for the mononuclear phagocyte growth factor, CSF-1. Cell 41, 665–676.

Syc-Mazurek, S. B. and Libby, R. T. (2019). Axon injury signaling and compartmentalized injury response in glaucoma. Prog. Retin. Eye Res. 73, 100769. doi:10.1016/j.preteyeres.2019.07.002

Takeda, A., Shinozaki, Y., Kashiwagi, K., Ohno, N., eto, K., Wake, H., Nabekura, J. and Koizumi, S. (2018). Microglia mediate non-cell-autonomous cell death of retinal ganglion cells. Glia 66, 2366–2384.

Todd, L., Squires, N., Suarez, L. and Fischer, A. J. (2016). Jak/Stat signaling regulates the proliferation and neurogenic potential of Müller glia-derived progenitor cells in the avian retina. Sci. Rep. 6, 35703. https://doi.org/10.1038/srep35703

Todd, L., Palazzo, I., Suarez, L., Liu, X., Volkov, L., Hoang, T. V., Campbell, W. A., Blackshaw, S., Quan, N. and Fischer, A. J. (2019). Reactive microglia and IL1β/IL-1R1-signaling mediate neuroprotection in excitotoxin-damaged mouse retina. J. Neuroinflammation 16, 118. https://doi.org/10.1186/s12974-019-1505-5

Tran, N. M., Shekhar, K., Whitney, I. E., Jacobi, A., Benhar, I., Hong, G., Yan, W., Adiconis, X., Arnold, M. E., Lee, J. M., et al. (2019). Single-Cell Profiles of Retinal Ganglion Cells Differing in Resilience to Injury Reveal Neuroprotective Genes. Neuron 104, 1039–1055.e12.

Tsarouchas, T. M., Wehner, D., Cavone, L., Munir, T., Keatinge, M., Lambertus, M., Underhill, A., Barrett, T., Kassapis, E., Ogryzko, N., et al. (2018). Dynamic control of proinflammatory cytokines Il-1β and Tnf-α by macrophages in zebrafish spinal cord regeneration. Nature Communications 9,4670 https://doi.org/10.1038/s41467-018-07036-w.

Uribe, R. A. and Gross, J. M. (2007). Immunohistochemistry on cryosections from embryonic and adult zebrafish eyes. CSH Protoc. 2007, db.prot4779.

Van Dyck, A., Bollaerts, I., Beckers, A., Vanhunsel, S., Glorian, N., van Houcke, J., van Ham, T. J., De Groef, L., Andries, L. and Moons, L. (2021). Müller glia-myeloid cell crosstalk accelerates optic nerve regeneration in the adult zebrafish. Glia. doi:10.1002/glia.23972

Villarino, A. V., Kanno, Y., Ferdinand, J. R. and O’Shea, J. J. (2015). Mechanisms of Jak/STAT signaling in immunity and disease. J. Immunol. 194, 21–27.

Warchol, M. E. (1999). Immune cytokines and dexamethasone influence sensory regeneration in the avian vestibular periphery. J. Neurocytol. 28, 889–900.

White, D. T., Sengupta, S., Saxena, M. T., Xu, Q., Hanes, J., Ding, D., Ji, H. and Mumm, J. S. (2017). Immunomodulation-accelerated neuronal regeneration following selective rod photoreceptor cell ablation in the zebrafish retina. Proc. Natl. Acad. Sci. U. S. A. 114, E3719–E3728.

Williams, P. A., Marsh-Armstrong, N. Howell, G. R. and Lasker/IRRF Initiative on Astrocytes and Glaucomatous Neurodegeneration Participants (2017). Neuroinflammation in glaucoma: A new opportunity. Exp. Eye Res. 157, 20–27.

Xie, Y., Tolmeijer, S., Oskam, J. M., Tonkens, T., Meijer, A. H. and Schaaf, M. J. M. (2019). Glucocorticoids inhibit macrophage differentiation towards a pro-inflammatory phenotype upon wounding without affecting their migration. Dis. Model. Mech. 12, dmm037887. doi: 10.1242/dmm.037887.

Zhang, C., Li, H., Liu, M.-G., Kawasaki, A., Fu, X.-Y., Barnstable, C. J. and ShaoMin Zhang, S. (2008). STAT3 activation protects retinal ganglion cell layer neurons in response to stress. Exp. Eye Res. 86, 991–997.

Zhao, X.-F., Wan, J., Powell, C., Ramachandran, R., Myers, M. G., Jr and Goldman, D. (2014). Leptin and IL-6 family cytokines synergize to stimulate Müller glia reprogramming and retina regeneration. Cell Rep. 9, 272–284.

Zou, S., Tian, C., Ge, S. and Hu, B. (2013). Neurogenesis of retinal ganglion cells is not essential to visual functional recovery after optic nerve injury in adult zebrafish. PLoS One 8, e57280.

